# Asymptomatic neonatal herpes simplex virus infection in mice leads to long-term cognitive impairment

**DOI:** 10.1101/2024.04.22.590596

**Authors:** A Dutton, CD Patel, SA Taylor, CR Garland, EM Turnbaugh, R Alers-Velazquez, J Mehrbach, KM Nautiyal, DA Leib

**Affiliations:** Department of Microbiology and Immunology, Geisel School of Medicine at Dartmouth, Lebanon, New Hampshire, USA; Guarini School of Graduate and Advanced Studies at Dartmouth, Hanover, New Hampshire, USA; Department of Psychology and Brain Science, Dartmouth College, Hanover, New Hampshire, USA

## Abstract

Neonatal herpes simplex virus (nHSV) is a devastating infection impacting approximately 14,000 newborns globally each year. Infection is associated with high neurologic morbidity and mortality, making early intervention and treatment critical. Clinical outcomes of symptomatic nHSV infections are well-studied, but little is known about the frequency of, or outcomes following, sub-clinical or asymptomatic nHSV. Given the ubiquitous nature of HSV infection and frequency of asymptomatic shedding in adults, subclinical infections are underreported, yet could contribute to long-term neurological damage. To assess potential neurological morbidity associated with subclinical nHSV infection, we developed a low-dose (100 PFU) HSV infection protocol in neonatal C57BL/6 mice. At this dose, HSV DNA was detected in the brain by PCR but was not associated with acute clinical symptoms. However, months after initial inoculation with 100 PFU of HSV, we observed impaired mouse performance on a range of cognitive and memory performance tasks. Memory impairment was induced by infection with either HSV-1 or HSV-2 wild-type viruses, but not by a viral mutant lacking the autophagy-modulating Beclin-binding domain of the neurovirulence gene γ34.5. Retroviral expression of wild type γ34.5 gene led to behavioral pathology in mice, suggesting that γ34.5 expression may be sufficient to cause cognitive impairment. Maternal immunization and HSV-specific antibody treatment prevented offspring from developing neurological sequelae following nHSV-1 infection. Altogether, these results support the idea that subclinical neonatal infections may lead to cognitive decline in adulthood, with possible profound implications for research on human neurodegenerative disorders such as Alzheimer’s Disease.

## Introduction

Herpes simplex virus (HSV) is a neurotropic DNA virus that establishes latency within sensory neurons^1,2,3^. Periodic reactivation of HSV may be asymptomatic, result in recurrent vesicular lesions, or lead to more severe infections including encephalitis and stromal keratitis^4^. These diverse disease manifestations usually follow anterograde transport of the virus within neuronal axons from sites of latency to sites of recrudescence^5^. Some of the worst outcomes follow primary neonatal infections in which pathological inflammation and direct damage can lead to severe and permanent neurological impairment and death^6,7^. Antiviral therapy decreases the frequency and severity of both symptomatic reactivation and neonatal disease, however, there is no cure or vaccine for HSV infection^8,9^. Current estimates suggest that two thirds of the global population under the age of 50 are infected with HSV, underscoring the large burden of both disease and potential asymptomatic transmission^10^.

Consistent with life-long persistence in sensory neurons and its ability to enter and cause inflammation within the brain, HSV has been implicated in a number of neurologic diseases, including Alzheimer’s disease (AD)^11–15^. AD is a chronic neurodegenerative disorder and the most common form of dementia in elderly populations^16^. AD and related dementias affect over 40 million people globally and are characterized by memory loss, anxiety, agitation, and depression^17,16^. These symptoms manifest from chronic inflammation, accumulation of neurotoxic proteins, and neuronal loss in the central nervous system (CNS)^18^. The chronic CNS inflammation induced by certain neurotropic pathogens supports the hypothesis that viral infection may play a role in the development of AD^19^. Post-mortem brains from individuals diagnosed with AD were discovered to have a high prevalence of HSV DNA, supporting a causal link^20^. The presence of HSV DNA in the CNS of individuals with AD, along with HSV’s propensity to cause neurologic morbidity, have implicated HSV in the pathogenesis of AD but the molecular mechanism is unknown^12,13^.

Mouse models have previously been utilized to investigate the connection between HSV and AD^21,22^. HSV-1-infected mice show CNS damage, inflammation, and an AD-like phenotype after multiple viral reactivations^21,22^. This model, however, requires a high, and likely super-physiological dose of HSV followed by multiple rounds of heat-induced reactivation to result in neurologic impairment. It is likely that human HSV infections result from significantly lower infectious doses, especially during asymptomatic shedding and transmission. Asymptomatic shedding, occurring during both acute infection and viral reactivation events, contributes to high rates of infection and exposure and makes infection in the human population difficult to track^23–26^. The majority of individuals who are seropositive for HSV are unaware of their infection^27,28^. We therefore worked to develop an infection model to study the relationship between low-dose, subclinical HSV infection and the development of cognitive decline.

To investigate the longitudinal implications of HSV infection, we utilized a neonatal model of infection. The neonatal period is an immunologically and neurologically vulnerable window of development^29–32^. Early life exposure to infectious or toxic insults can perturb development and lead to neurologic and psychiatric disease in later life^33,32^. While serious acute cases of neonatal HSV (nHSV) are well studied, the burden of asymptomatic, life-long nHSV infections remains unknown. Low rates of testing during pregnancy^34^, and the fact that nHSV is not diagnosed until symptoms appear^35–37^, jointly contribute to the possibility of asymptomatic nHSV cases going undiagnosed and untreated. Given the high prevalence of HSV infection in the adult population, asymptomatic HSV infection is likely an unrecognized source of morbidity that may have lifelong implications for infected neonates. In this study, we aimed to characterize a novel asymptomatic nHSV infection model and investigate the role of neonatal viral infection in cognitive decline in adulthood. Findings from this work adds support to studies suggesting a link between aging and early life exposure to pathogens, and further supports the hypothesis of a connection between HSV infection and AD.

Low-dose neonatal HSV (nHSV) infection causes long-lasting, anxiety-like behavior in mice^38,39^. To improve our understanding of how asymptomatic neonatal infections may contribute to long-term neurological disease, we examined the effect of subclinical nHSV infection on adult memory loss and cognitive decline. Low-dose nHSV-1 infection led to memory-loss and decreased behavioral flexibility several months after initial infection. We determined that the Beclin-1-binding function of HSV γ34.5 was required for this activity, and that retroviral expression of γ34.5 was sufficient to cause cognitive decline. The behavioral sequelae induced by nHSV were prevented if mice were born to HSV-seropositive or immunized dams, demonstrating that maternally derived HSV-specific antibody was protective against neonatal impairment. Importantly, in this era of vaccine hesitancy^40,41^, we showed that vaccination alone did not cause cognitive decline. Taken together, the findings in this study provide compelling evidence that HSV-1 can cause AD-like cognitive decline in mice, specifically through neurovirulence conferred by the Beclin-1-binding domain of γ34.5 during the neonatal period.

## Results

### Characterization of a low-dose neonatal HSV infection model

To model the impact of neonatal exposure to small amounts of HSV, we titrated viral inocula with a goal of achieving CNS infection without acute symptoms (Fig. 1A). On day 1 of life (P1), litters of mice were intranasally (i.n.) infected with escalating doses of strain17 (st17) HSV-1 or mock-infected with Vero cell lysate. 5 days post-infection (dpi), brains of mice were collected to determine viral burden. Infection with 100 plaque-forming units (PFU) st17 HSV-1 resulted in 90% survival while 40% of pups survived following infection with 1000 PFU (Fig. 1B). Notably, the mice that survived infection with 100 PFU of HSV showed no clinical symptoms of acute infection. To broadly determine the extent of HSV spread in the CNS following i.n. infection, brains of 1000 PFU infected pups were sectioned and assessed by immunofluorescence 5 dpi (Fig.1C, top). Significant fluorescence, indicating presence of HSV antigen, was detected in the frontal cortex, mid-brain, and brainstem, indicating that i.n. infection leads to a detectable, yet localized infection of the CNS (Fig. 1C, bottom). Virus-specific staining in brains of mice infected with 100 PFU was undetectable by immunofluorescence, but significant genome load in the midbrain at 5 dpi was detectable by quantitative DNA PCR (Fig. 1D). As expected, higher viral inoculum resulted in greater genome loads. Therefore, we determined that 100 PFU HSV-1 was the optimum inoculum to establish detectable CNS infection without causing significant mortality or morbidity at the neonatal timepoint.

**Figure 1:**
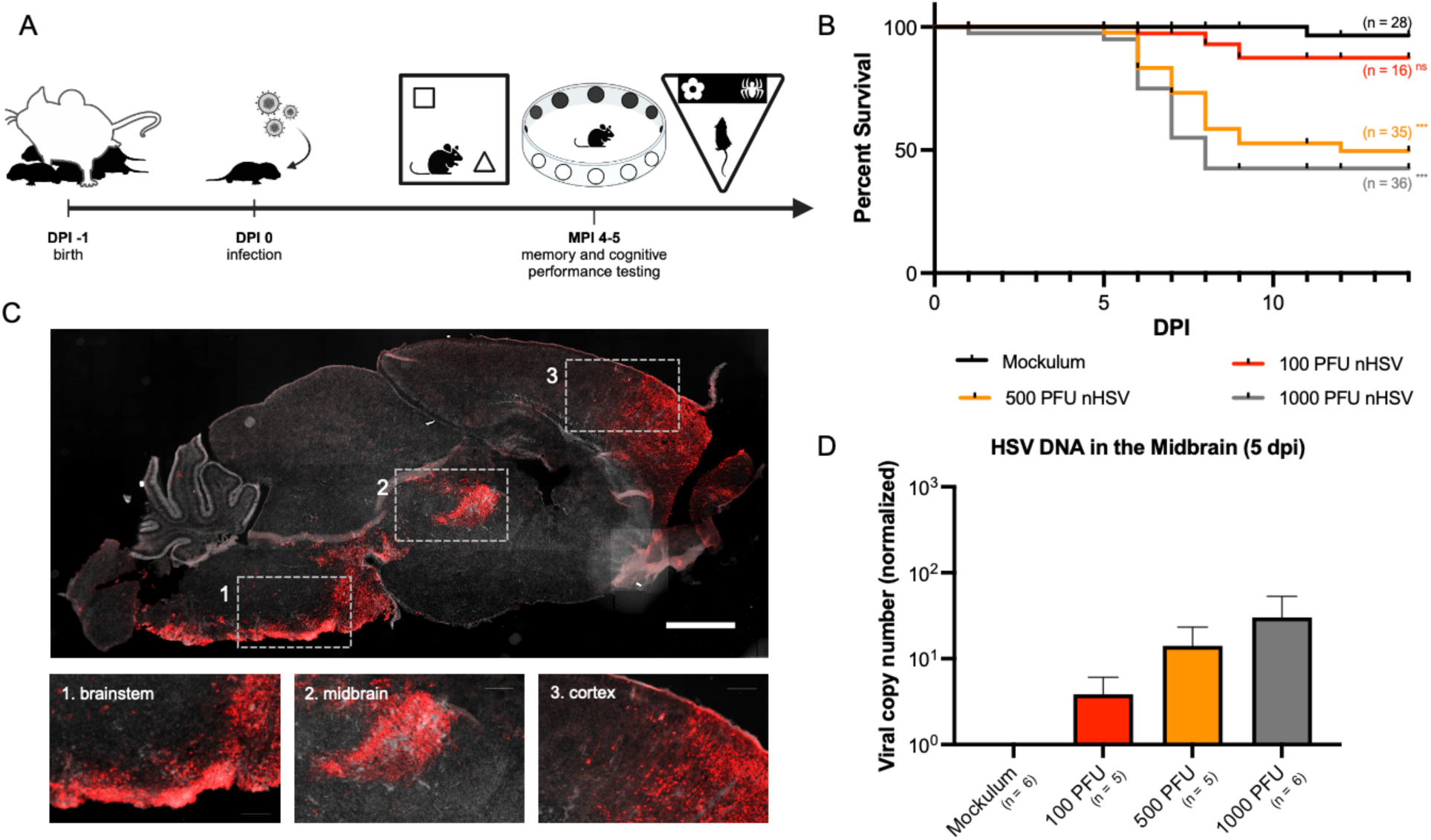
Characterization of a low-dose neonatal HSV infection model. (A) Timeline of infection and behavioral assessment. One-day-old C57BL/6 mice were intranasally (i.n.) infected with escalating doses of st17 HSV-1 or virus-free “mockulum”. At 5 mpi, nHSV and mockulum-treated mice were assayed for memory and cognitive performance deficits using the novel object recognition (NOR) task, the novel object location (NOL) task, the modified Barnes maze (MBM), and the different paired associative learning (dPAL) task. (B) Survival of mice intranasally infected with mockulum, 100 PFU, 500 PFU, or 1000 PFU HSV-1. (C) Representative brain slice of a 5 dpi 1000 PFU HSV-1-infected mouse stained with DAPI and antibody against HSV glycoprotein C (gC). Scale bar = 1mm. (D) Statistical significance was determined by comparing infected groups to mock-infected animals using log-rank (Mantel-Cox) tests (A) and multiple unpaired t-tests (D). ***p<0.001, ns = not significant. Error bars represent standard error (SEM).

### Low-dose nHSV infection causes cognitive and memory performance deficits in adulthood

Mice infected with nHSV-1 demonstrate anxiety-like behavior in the open field fest (OFT) 5 weeks after infection^38^. Given the possible link between HSV infection and Alzheimer’s disease^11–13^, we tested whether neonatal infection with 100 PFU HSV-1 would also lead to memory loss in adult mice. To test this, we used the novel object recognition (NOR) task to assess recognition memory at 4-5 months post infection (mpi). The NOR task is derived from the visual paired-comparison paradigm used in human studies to assess recognition memory in the context of disease and/or aging^42^. In the NOR task, on day 1 (familiarization phase), mice are trained to recognize two identical objects placed in an open enclosure for 10 minutes. On day 2 (testing phase), one object is replaced by a novel object and interaction time with each object is recorded (Fig. 2A top). Naïve mice spend more time exploring the novel object, indicating intact recognition memory of the familiar object^43^. At 5 mpi, mock-infected mice spent more time with the novel object, demonstrating intact novel-seeking behavior and recognition memory. nHSV-infected mice, however, could not discriminate between novel and familiar objects and spent approximately equal time with both (Fig. 2A bottom). The average discrimination index (DI; a metric of relative preference^44,45^) for mock-infected mice was significantly higher than the average discrimination index for nHSV-infected animals (t(37)=2.233, p = 0.0317), indicating that nHSV infection lead to impaired novel object recognition at 5 months of age.

**Figure 2:**
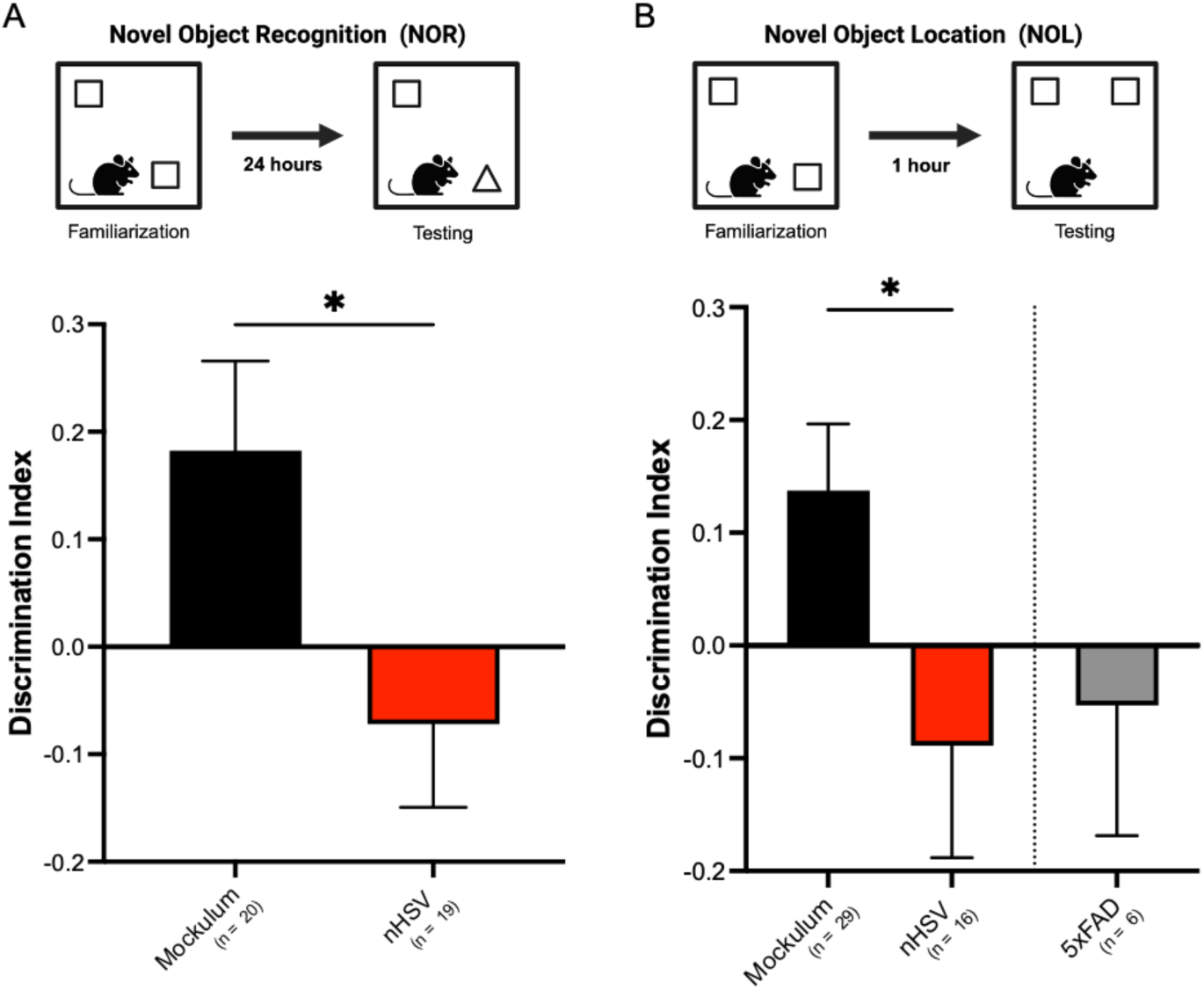
Low-dose nHSV infection leads to impaired novelty recognition in adult mice. (A) Novel object recognition (NOR) task performance of mockulum-treated (n=20) and st17 nHSV- infected (n=19) mice at 5 mpi. Data was collected and quantified as described in materials and methods. Bar graphs show the mean discrimination index for the novel object. (B) Novel object location (NOL) recognition performance of 5 mpi mockulum-treated mice (n=29), 5 mpi st17 nHSV- infected mice (n=16), and 7-month-old uninfected female 5xFAD mice (n=6). Data was recorded and quantified as described in materials and methods. Bar graphs show the mean discrimination index for the object in a novel location. Discrimination index = (Time_novel_ - Time_familiar_)/Time_total_. Statistical significance was determined by unpaired t-tests. *p<0.05. Error bars represent standard error (SEM).

To test infected mice’s spatial memory, we used the novel object location (NOL) task, described previously^45,46^. Spatial memory impairment is one of the hallmark deficits associated with AD, shared by over 60% of those with a diagnosis^47,48^. This task is identical to the NOR task, except an object is moved to a novel location during the testing phase rather than replaced with a novel object.

Additionally, the time between training and testing was reduced to 1 hour rather than 24 hours to assess short-term spatial recognition memory (Fig. 2B top). Mock-infected mice spent more time with the object in its novel location than nHSV-infected mice, demonstrating that spatial memory recognition is impaired following nHSV infection (Fig. 2B bottom). Interestingly, this NOL impairment mirrors our observations of 7-month-old female 5xFAD model mice performance on the NOL task (Fig. 2B). 5xFAD mice express 5 familial AD mutations resulting in overexpression of APP and PSEN1 and exhibit early onset amyloid pathology and cognitive performance deficits^49,50^. Like nHSV-infected mice, 7-month-old female 5xFAD mice did not show preference for the object in its novel location, indicating shared spatial memory impairment.

Multiple brain regions are involved in recognition memory processing, including the perirhinal cortex, the medial prefrontal cortex, the hippocampus, and the medial dorsal thalamus^51^. To further define and map memory impairment associated with nHSV infection, we used the modified Barnes maze (MBM), and the different paired-associate learning (dPAL) paradigm. The MBM is a spatial learning task in which mice use visual orientation cues to navigate a circular enclosure and find an escape hole^52,53^. A reversal phase allows for assessment of reversal learning and behavioral flexibility encoded by the prefrontal cortex^54^ (PFC; Fig. 3A). Mock-infected and nHSV-infected mice performed similarly during the training phase of the assay (Fig. 3B). In contrast, during the reversal phase, nHSV-infected mice took longer to find the escape hole (Fig. 3C). On day 1 of reversal testing, nHSV-infected mice traveled significantly more distance (t(8)=4.225, p=0.0029), and made significantly more errors per trial (t(8)=2.889, p=0.0202) than mock-infected mice (Fig. 3D-E). Importantly, of the errors made, the majority were committed in the quadrant of the maze that held the original exit hole during training. An increase in error rate in this quadrant is indicative of perseveration and impaired behavioral flexibility, consistent with nHSV-induced PFC impairment^55^.

**Figure 3:**
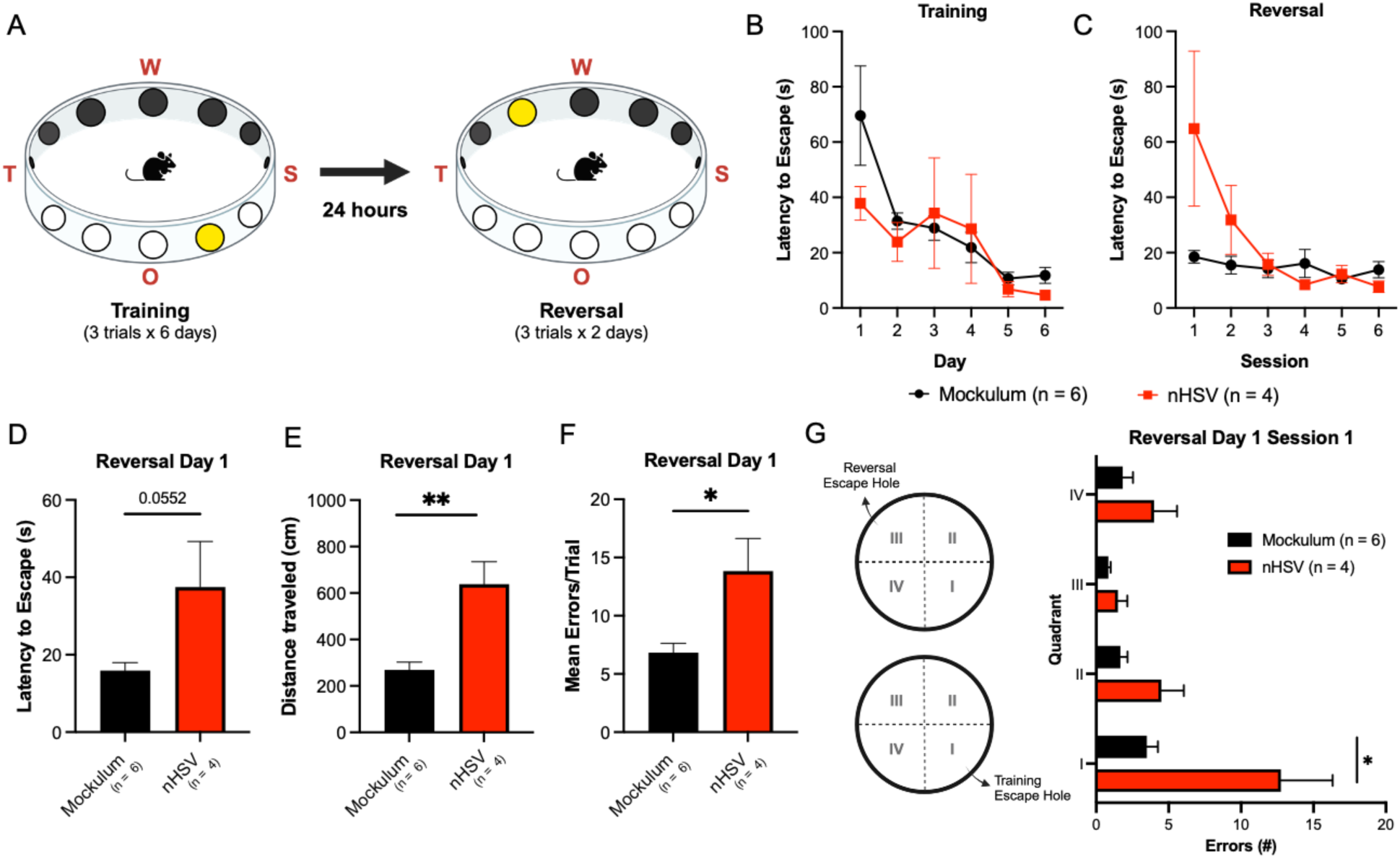
Low-dose nHSV infection leads to impaired behavioral flexibility in adult mice. (A) Over the course of six days of 3 trials per day, mice were trained in a modified Barnes maze (MBM) arena to locate, and exit through, an escape hole using visual cues (“W”, “S”, “O”, and “T”) for orientation. Following training, the location of the exit hole was reversed, and mice were tested for their ability to adapt and locate the reversal phase exit. (B) Mean latency to escape per training day during the training phase. (C) Mean latency to escape per session during the reversal testing phase. (D) Mean latency to escape on the first day of reversal testing. (E) Mean distance traveled in centimeters (cm) during the first day of reversal testing. (F) Mean errors per trials committed during the first day of reversal testing. (G) Quadrant designations for the location of the escape hole during training (I) and reversal (III) phases (left). Mean number of errors per quadrant committed during the first reversal session on day one of reversal testing (right). Statistical significance was determined by unpaired t-tests (D-G). *p<0.05. Error bars represent standard error (SEM).

To probe hippocampal-dependent behavior^56^, we tested mice using the dPAL task on a touch screen behavioral operant system^56^. A similar version of this task is used to assess memory cognition in AD patients^57^. In mice, dorsal hippocampal lesions impair ability to perform well on this task but do not disrupt task acquisition^56^. Therefore, dPAL is a useful behavioral assay to assess hippocampal performance deficits. Following pre-task training, we tested mock-infected and nHSV-infected mice at 5 mpi. Each day, mice completed up to 36 trials in which a correct response was noted as an interaction with the shape (spider, flower, or plane) in its correct location (Fig. 4A). Both groups exhibited evidence of learning over time and reached an average of 75% correct responses by the final day of testing (day 40). The nHSV-infected group, however, showed a reduced proportion of correct responses compared to the mock-infected group during initial days of testing. In the initial 5 blocks of testing, there was a significant effect of progressive testing (F(4,175)=9.300, p<0.0001) and of infection status (F(1,44)=7.299, p<0.01; Fig. 4B) on trial performance (% correct). In addition, nHSV-infected mice completed a greater number of correction trials than mockulum-treated mice. In the initial 5 blocks of testing, there was a significant effect of progressive testing (F(3.426, 150.7)=25.5, p<0.0001) and infection status (F(1,44)= 6.956, p<0.05; Fig. 4C) on the number of correction trials made. Together, these trends indicate that both groups of mice improved in dPAL performance over time. That said, in the initial 5 blocks of testing, nHSV-infected mice had a lower performance (% correct) and completed a greater number of correction trials than mockulum-treated mice. On the first day of dPAL testing, nHSV- infected mice scored significantly lower on the task than mock-infected mice and completed significantly more correction trials (Fig. 4E-F). Importantly, both groups completed approximately the same number of trials during the first hour session indicating that performance deficits were not due to differences in exposure to the paradigm (Fig. 4G). By the fifth block of testing, nHSV-infected mice performed similarly to mock-infected mice, perhaps indicating that nHSV infection has a greater impact on short-term spatial memory than long-term memory (Fig. 4 B-C). Taken together, mouse performance on the NOR, NOL, MBM, and dPAL tasks demonstrates that neonatal infection with HSV leads to changes in memory and cognitive behavior similar to those seen in AD^58^.

**Figure 4:**
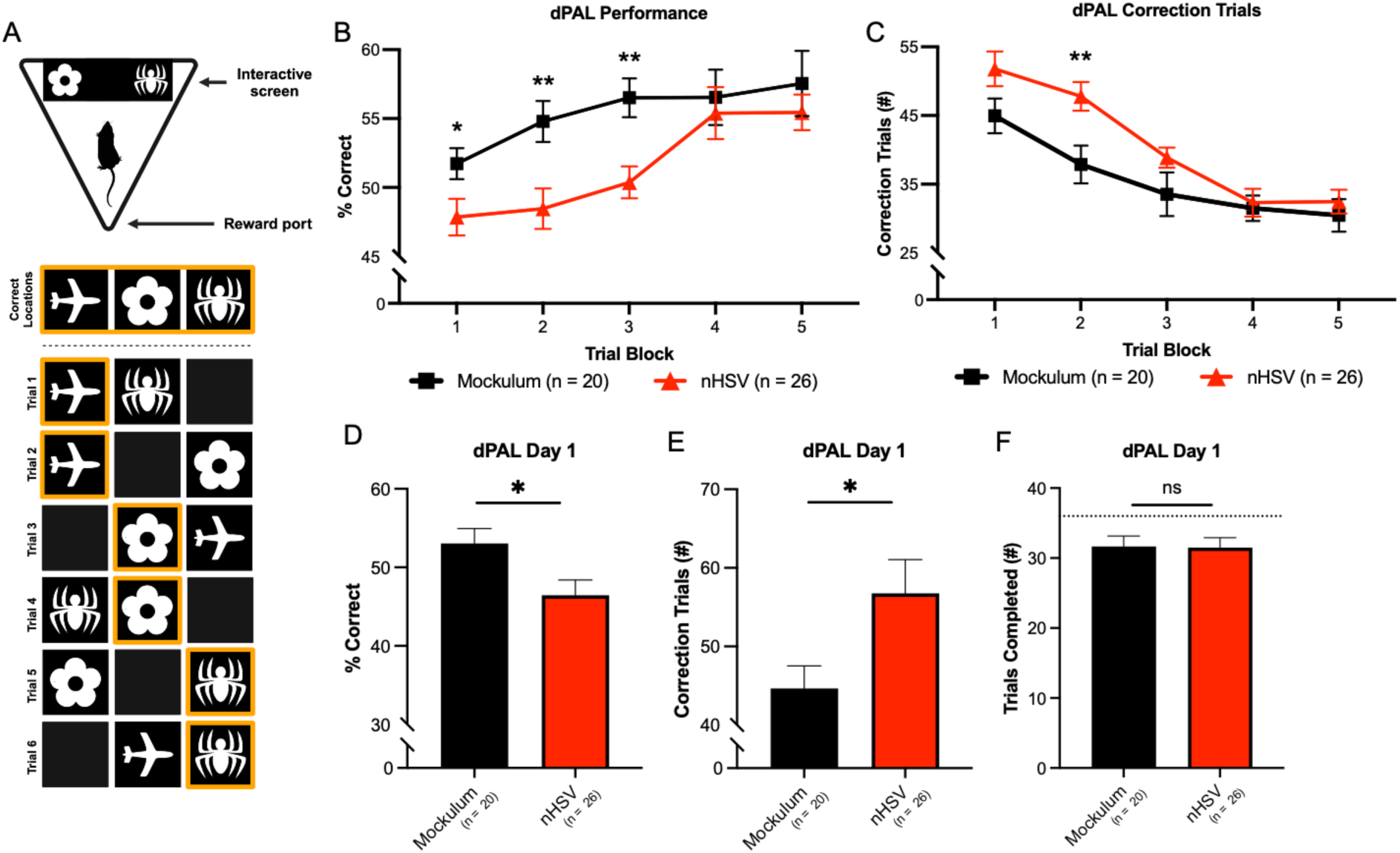
Low-dose nHSV infection leads to impaired associative memory in adult mice. (A) Cartoon of a mouse completing the paired associative learning (dPAL) paradigm in a Bussey Box chamber (top), and a diagram representing all possible combinations of test stimuli and correct responses (orange; bottom). (B) Average performance (% correct on first attempt) of mock-infected (20) and st17 nHSV-infected (26) mice during the first 5 trial blocks. 1 block = 3 testing sessions = 108 trials completed across 3 days. Multiple unpaired t-tests show significant differences between infection groups on blocks 1 (p<0.05), 2 (p<0.01), and 3 (p<0.01). (C) Average number of correction trials per testing block during the first 5 blocks. Multiple unpaired t-tests show significant differences between infection groups on block 2 (p<0.01). (D) Average percent correct on day 1 of dPAL testing. (E) Average number of correction trials on day 1 of dPAL testing. (F) Average number of total trials completed on day 1 of dPAL testing. Statistical significance was determined by unpaired t-tests. *p<0.05. Error bars represent standard error (SEM).

To better characterize our nHSV model, we examined the ability of other HSV strains to induce behavioral morbidity. Given that HSV-2 predominates as the causative agent of nHSV in many parts of the world^59,10^, we wished to examine whether nHSV-2 infection could cause learning and memory deficits as shown for HSV-1. We adjusted the i.n. dose of HSV-2 strain 333 for neonatal mice to 75 PFU because HSV-2 is more virulent than HSV-1 and a dose reduction was required to attain an asymptomatic infection^60,61^. The mice neonatally infected with HSV-2 showed no preference for the familiar object in the NOR task at 4-5 mpi indicating that, consistent with 100 PFU st17 HSV-1 neonatal infection, there was adult memory impairment associated with neonatal HSV-2 infection (Fig. 5A).

**Figure 5:**
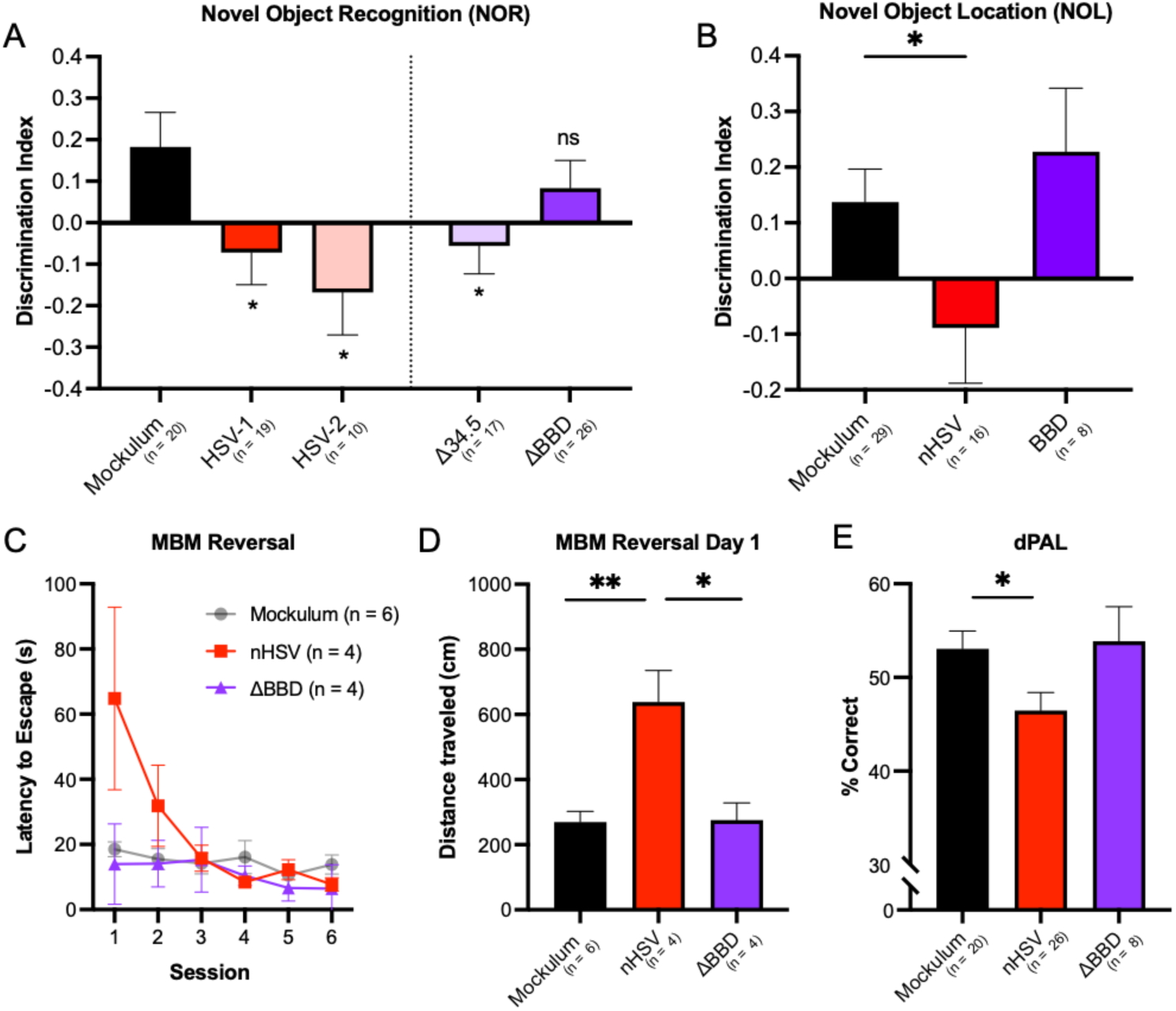
nHSV-induced behavioral morbidity is dependent on the HSV γ34.5 Beclin-1-binding domain. (A) Average NOR task performance of neonatally infected mice at 5 mpi. Data are shown as average discrimination index for novel object during the testing phase of the NOR task. Statistical significance given as compared to mockulum-treated mice. (B) Average discrimination index in the NOL task at 5 mpi. (C) Mean latency to escape per trial in the MBM task during the reversal testing phase at 5 mpi. (D) Mean distance traveled in cm during the first day of reversal testing. (E) Average percent correct on the dPAL task on day 1 of testing. Statistical significance determined by unpaired t-tests. *p<0.05, **p<0.01. Error bars represent standard error (SEM).

### nHSV-1 induced behavioral changes are dependent on HSV ICP 34.5 Beclin-1-binding

Given the possible links between AD, neurotropic virus infections, and autophagy dysregulation^62–64^, we wished to assess the role of γ34.5 in the induction of behavioral morbidity in our neonatal model. HSV γ34.5 encodes a multifunctional protein that interferes with host-mediated shutoff of protein synthesis through PKR, type I interferon production, and host-cell autophagy^65–67^. We therefore neonatally infected mice with HSV-1 mutants Δ34.5 (γ34.5 null mutant), or ΔBBD (a mutant lacking the 20aa Beclin-binding domain (BBD). γ34.5 reverses PKR-mediated translational arrest in infected cells while BBD confers neurovirulence more specifically through inhibition of host-cell autophagy and control of CD4+ T-cell responses^66–68^. Mice neonatally infected with Δ34.5 demonstrated impaired object recognition in the NOR task, similar to mice infected with WT HSV (Fig. 5A). Mice neonatally infected with ΔBBD, however, demonstrated intact object recognition memory on NOR, spending significantly more time with the novel object than the familiar object, similar to mock-infected mice at 5 mpi (Fig. 5A).

To add to our findings from the NOR task, we repeated the NOL, MBM, and dPAL tasks on mice neonatally infected with ΔBBD. On the NOL task, adult mice that had been infected as neonates with 100 PFU ΔBBD spent significantly more time with the object in a novel location than in the familiar location, similar to mock-infected mice (Fig. 5B). On the reversal phase of MBM, ΔBBD infected mice performed similarly to mock-infected mice. They took less time to find the escape hole than WT nHSV- infected mice and traveled significantly less distance to find the exit (Fig. 5C-D). Finally, ΔBBD-infected mice performed significantly better on day 1 of dPAL testing than WT nHSV-infected mice (Fig. 5E).

These data suggested that memory impairment is dependent on the presence of ICP34.5 BBD, but the absence of the additional functions of γ34.5 renders the virus capable of inducing memory loss.

### Expression of HSV-1 ICP34.5 is sufficient to cause anxiety and memory loss

To further examine the relationship between ICP34.5 Beclin-1 binding and memory impairment, we utilized lentiviral vectors (LV) to express full length (LV 34.5) and BBD-deleted (LV ΔBBD) ICP34.5 independently of HSV-1 infection. LV 34.5 or LV ΔBBD were administered intranasally to neonatal mice (10,000 transducing units) using empty LV vector as an additional control (Fig. 6). At 4 mpi, in the NOR task, mice infected with LV 34.5 spent less time exploring the novel object, while LV ΔBBD-infected mice demonstrated intact object recognition spending more time with the novel object, similar to empty vector-infected control mice (Fig. 6). LV ΔBBD transduced mice therefore had intact object recognition memory in contrast to mice transduced with a vector expressing full length ICP34.5. These findings are consistent with the hypothesis that modulation of Beclin-1 induces behavioral sequelae in mice.

**Figure 6:**
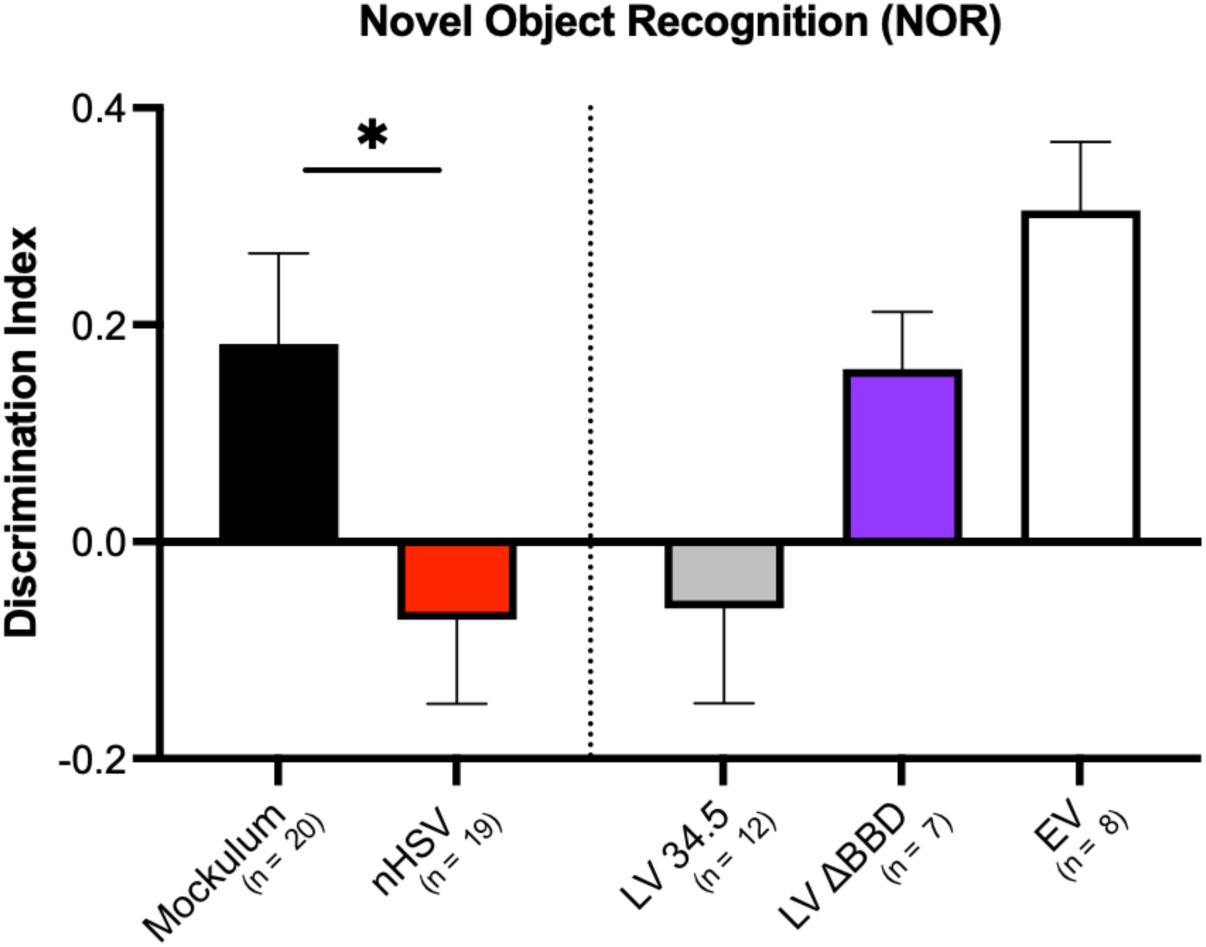
Transduction of neonatal mice with lentiviruses expressing full length HSV-1 γ34.5, but not Λ1BBD, causes memory impairment in adulthood. Average NOR task performance of nHSV-infected and neonatally lentiviral-transduced mice at 5 mpi. Mice were either infected i.n. with HSV-1 or transduced i.n. with lentiviruses expressing full length γ34.5, Λ1BBD, or an empty lentivirus. Data are shown as average discrimination index for novel object during the testing phase of the NOR task. Discrimination index = (Time_novel_ - Time_familiar_)/Time_total_. Statistical significance was determined by unpaired t-tests. *p<0.05. Error bars represent standard error (SEM).

Importantly, this effect can be observed even in the absence of HSV-1 infection, demonstrating that ICP34.5 is sufficient to cause memory loss following neonatal infection.

### HSV-specific maternal immunity protects against nHSV-1 induced behavioral morbidity

Maternal immunity is critical to the outcome of nHSV morbidity and mortality in mice^38,69^. Both passive and active immunization confers HSV-specific antibody protection in dams that lowers viral titers and anxiety behavior in adult offspring neonatally challenged with HSV^38,69^. Given this, we determined the ability of maternally derived Ab to provide lasting protection against memory loss and cognitive impairment. Neonatally HSV-challenged offspring of mice from seropositive dams were tested using the NOR task at 11-12 mpi (Fig. 7A). To induce seropositivity, dams were either (1) latently HSV-infected via the ocular route, prior to parturition, (2) treated with HSV-specific IgG, or (3) vaccinated with live-attenuated virus (*dl*5-29) via intramuscular injection. Control dams were given mockulum or control IgG and were therefore seronegative. Neonatally HSV-challenged offspring of seronegative dams spent less time with the novel object than with the familiar object while offspring of seropositive dams exhibited intact object recognition memory behavior (Fig. 7B). To assess the risks of live vaccine itself (*d*l5-29 in this model) conferring memory impairment, we i.n. infected neonatal mice with 100 PFU of replication-deficient HSV vaccine strain *dl*5-29 and followed up with NOR testing at 4 mpi. At 4 mpi, mice i.n. infected with *dl*5-29 did not display object recognition memory impairment on the NOR task and exhibited performance comparable to mock-infected mice (Fig. 7C). This lends support for its safety as a vaccine, and together, these data indicate that maternal immunity is a critical determinant of protection against HSV-induced memory impairment in offspring.

**Figure 7:**
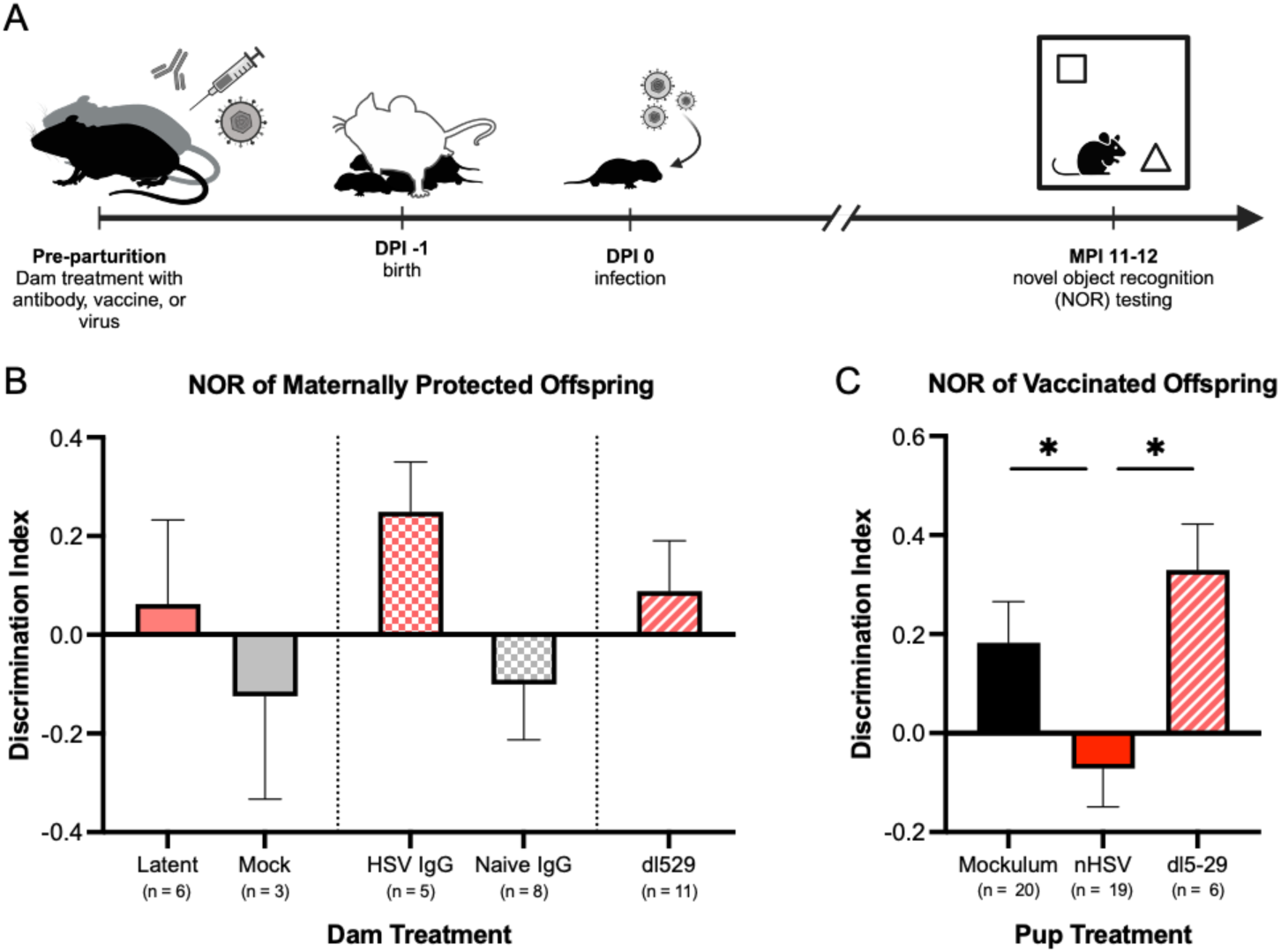
Maternally derived antibody protects against nHSV-induced behavioral morbidity. (A) Timeline of dam treatment, neonatal infection, and behavioral assessment using the NOR task. (B) Average NOR performance of offspring of dams treated with latent HSV infection, mockulum, HSV IgG, naïve IgG, or vaccination with replication deficient HSV strain dl529. (C) Average NOR performance at 5 mpi of mice neonatally infected with replication deficient HSV vaccine strain dl526. Data are shown as average discrimination index for novel object during the testing phase of the NOR task. Discrimination index = (Time_novel_ - Time_familiar_)/Time_total_. Statistical significance determined by unpaired t-tests. *p<0.05. Error bars represent standard error (SEM).

## Discussion

Herpes simplex viruses are a highly prevalent part of the human virome and the lifelong presence of HSV in the nervous system likely impacts neuroimmune balance and homeostasis. HSV CNS infections are common causes of encephalitis and neurological dysfunction^70–73^, but the contribution of HSV infections to pathological changes to memory and cognition remains underexplored. In this study, we developed a mouse model of neonatal asymptomatic infection and demonstrated that nHSV-1 and nHSV-2 causes memory loss and behavioral changes many months after initial infection. Importantly, we have shown that these morbidities occur in the absence of clinically observable symptoms, such that even asymptomatic infection may lead to cognitive impairment. We further demonstrate that the BBD of γ34.5 may be required for the development of memory loss. These studies provide evidence for a possible link between viral infection, autophagy, and neurologic dysfunction, in addition to a possible mechanism for the development of HSV-induced cognitive decline.

Our study results are consistent with previous studies that showed cognitive decline, hippocampal neurodegeneration, and neuroinflammation in mice with recurrent HSV-1 infections^13,21,22^. Our model, however, differs in two important ways: (1) We detected behavioral and neuropathologic changes without induction of reactivation with thermal stress, suggesting that induced reactivation may not be necessary for cognitive decline in our model; (2) The dose used in previous studies was 10,000-fold greater than the dose used in this work^21^. This allowed us to examine the effect of a subclinical, asymptomatic infections on cognition without the confounding effects of significant inflammation and damage to the CNS. It is possible that our mice experienced low-level viral reactivation in the absence of thermal stress. Experiments utilizing acyclovir during the period between infection and cognitive testing may help elucidate this possibility. The very low HSV inoculum needed to induce memory impairment, however, suggests that virally induced cognitive decline may not depend on reactivation events in our model. Rather, the exquisite sensitivity of neonatal mice to infection likely predisposes the CNS to local inflammation and cognitive decline that is clinically inapparent until later in life. Our hypothesis is that perturbation during the neonatal period can affect neurodevelopment at a period in life during which the brain is rapidly changing. Many known maternal and fetal insults are accompanied by long-term behavioral alterations in offspring^29,32^. Prenatal alcohol exposure, for example, causes changes in neuronal migration in the prefrontal cortex leading to behavioral deficits^74^. It is possible that HSV invasion of the CNS during development, as well as the ensuing immune response, can shift neuronal differentiation or migration, leading to lifelong changes in the brain. If this model is translatable to human infections, the public health implications are profound.

We showed that HSV-associated memory loss is associated with a key activity of γ34.5, namely its ability to interact with Beclin-1 and modulate autophagy through the BBD. The BBD of HSV-1 is a 20aa domain of ICP34.5 which confers neurovirulence through control of CD4+ T-cell responses and inhibition of host-cell autophagy^66,67^. This is consistent with inflammation and autophagy being important determinants of behavioral morbidity following HSV infection. One paradox, however, is that while infection with ΔBBD did not induce cognitive morbidity, infection with the null mutant Δ34.5 induced memory loss in the NOR task. A possible explanation is that the neuro-attenuation of Δ34.5 derives from the lack of the multiple domains that inhibit immune response pathways, such as STING, antigen presentation, dendritic cell maturation, and IRF3 activation, that could lead to increased innate responses and inflammation^66^. The consequences of this inflammation may lead to neurological damage that masks the added inability of this mutant virus to control autophagy.

The mechanism by which autophagy interference by HSV could cause neurodegenerative pathology may be multifactorial. HSV seeds amyloid peptide in the brain^75^ and reactivation and subsequent Aβ production may lead to the pathological accumulations associated with AD, as well as the behavioral changes seen in our model. These accumulations may be accelerated if HSV-1 actively blocks host autophagy responses through γ34.5, either acutely or during a reactivation event. Of note, Beclin-1 is a target of multiple common human viruses. The HIV NEF^76^, HCMV TRS1^77^, and HFMD 2C virus^78^ all bind Beclin-1 to block autophagosome formation and maturation. It would be compelling to study whether seropositivity of these viruses is in any way associated with AD, as has been shown for HSV.

Our work is the first, to our knowledge, to show memory impairment following low-dose neonatal infection. Given that 70% of the population sheds HSV-1 asymptomatically at least one time per month^25^, there is a significant possibility of low level neonatal exposure. Therefore, our model captures a population that is not represented by previous studies on the link between infection and neurodegeneration. Although the host response following low-dose infection remains to be fully explored, our work elucidates the potential for subclinical, asymptomatic infections to cause long term changes to behavior and cognitive decline.

Impaired neurogenesis, neuronal loss, and accumulation of misfolded protein are additional hallmarks of AD which we have yet to explore. Our previous work has shown that nHSV induces anxiety-like behavior^38^. The work described here characterizes memory loss associated nHSV using NOR, NOL, the MBM, and dPAL tests. On the NOR and NOL, infected mice showed impaired object and spatial novelty recognition, perhaps indicating memory deficit. On the MBM, infected mice showed impaired performance on the reversal phase of testing, indicating impairment in PFC-mediated behavioral flexibility^55^. On dPAL, infected mice demonstrated impaired hippocampal learning and memory relative to mock-infected mice. Together, performance on these behavioral tasks point to areas of the brain that may be particularly vulnerable to neonatal HSV infection or consequential neurodegenerative processes. Ongoing work in our lab aims to characterize the lifelong neuroanatomical consequences of neonatal HSV exposure.

Our lab previously showed that maternal HSV-specific antibody is transplacentally transferred to the nervous system of the fetus in humans and mice^69^, and we demonstrated that maternal antibody prevents nHSV-induced cognitive impairments^38^. Maternal HSV-specific antibody arising from latent infection, immunization with an HSV-2 replication-deficient virus (*dl*529), or HSV-specific IgG was sufficient to protect against cognitive impairment. These maternal antibodies potently neutralize and protect against disseminated neonatal infection and suggest that the prevention of HSV viral entry precludes development of HSV-induced long-term memory loss. Maternal immunity, therefore, in the context of neonatal HSV infection, is more crucial than formerly thought, and may provide the most effective protection against nHSV and its sequelae. If nHSV infections potentiate memory loss, the safety of live attenuated vaccine strains must be carefully considered. Importantly, the HSV-2 *dl*5-29 replication-deficient vaccine strain did not lead to memory impairment after neonatal infection, reinforcing the idea that this represents both an efficacious and safe vaccine candidate. Beyond vaccines, many companies are developing HSV-based oncolytic vectors for neuroblastoma and CNS gene delivery^79,80^. It follows from this study that these vectors should be analyzed for their ability to cause CNS damage and cognitive issues in the long term. Finally, the effects of prior HSV infection must be explored. The HSV-1 seropositive population is over 60% and so it is important to consider both therapeutic and prophylactic approaches^59^. The HSV antiviral valacyclovir, is currently in Phase 2 clinical trials for treatment of AD by the National Institute on Aging (NCT03282916). In the double-blind trial, 130 patients with mild AD will be given valacyclovir or placebo daily for 78 weeks and monitored for cognitive function and accumulation of amyloid and tau protein. Results from this trial have the potential to inform our understanding of the relationship between viral Infection and AD, as well as contribute to novel therapeutic approaches for the treatment of AD.

A direct link between HSV and neurodegeneration has remained elusive since the early 1990s when the association was first proposed^12^. Previous research linking HSV with AD points to three hypotheses for an infectious etiology of AD. First, multiple reactivations of latent virus could produce chronic brain disease and progressive injury^21^. Second, that acute infection creates a pro-inflammatory response that exacerbates chronic neurodegeneration^81^. Third, that viral infection seeds beta amyloid peptide (Aβ) in the brain^75^. Our current work cannot exclude any of these non-mutually exclusive hypotheses, but this neonatal model will allow us and others to design future experiments to better define the mechanisms by which HSV infection leads to cognitive decline.

## Materials and Methods

### Cells and viruses

The HSV-1 strains used in this study were HSV-1 strain 17syn+^82^, HSV-2 strain 333, 17Δ34.5^83^, and 17ΔBBD (Δ68-87)^66^. Immunization studies were carried out using HSV-2 *dl*5-29, which lacks UL5 and UL29 and is derived from HSV-2 186 syn+^84^, generously provided by David Knipe (Harvard University).

Lentiviral constructs were made from the FUC-Cas9 plasmid (generously provided by Dr. Bryan Luikart, Dartmouth)^85^. Cas9 was removed by digestion with EcoRI and replaced with HA-tagged ICP34.5, wildtype or ΔBBD. The resulting plasmid contained mCherry and the ICP34.5 variant under the ubiquitin promoter linked by the T2A self-cleaving peptide. To produce lentiviruses, we transfected the ICP34.5 transfer plasmid, pCMV-VSV-G^86^, and pCMV-d8.9 into Lenti-X 293T cells using jetPRIME (Polyplus), then harvested supernatant and concentrated by sucrose gradient centrifugation^87^. Viruses were titered by counting fluorescent cells after transduction.

Virus stock preparation and the plaque assay were performed using Vero cells as described previously^88^. Vero cells were cultured in Dulbecco’s modified Eagle’s medium supplemented with 5% fetal bovine serum, 250 U/ml penicillin, and 250 µg/ml streptomycin.

### Mice and animal procedures

All procedures were performed in accordance with federal and university policies. The mice used in this study were C57BL/6 (B6) and 5xFAD (B6.Cg-Tg(APPSwFlLon,PSEN1*M146L*L286V)6799Vas/Mmjax; Strain #034848-JAX) mice. The B6 mice were purchased from The Jackson Laboratory (Bar Harbor, ME) and the 5xFAD mice were provided by Dr. Charles Sentman, Dartmouth) and bred in the barrier facility in the Center for Comparative Medicine and Research at the Geisel School of Medicine at Dartmouth.

### Viral challenge

Neonatal mice postnatal day 1-2 (P1-2) were infected intranasally (i.n.) with 10^2^-10^3^ PFU HSV or 10,000 transducing units of lentivirus in a volume of 5μl under 1% isoflurane anesthesia. Pups were monitored for weight loss and survival. Endpoints for survival studies were defined as excessive morbidity (hunching, spasms, or paralysis) and/or >10% weight loss (Fig. 1B). For maternal infections, adult mice were corneally infected as previously described^88^. Briefly, following corneal scarification, 2 x 10^5^ PFU st17 HSV-1 in 5μl was placed on the surface of each cornea of the mouse while under isoflurane anesthesia.

### Immunization

For the *dl*5-29 immunization, eight-week-old B6 female mice were immunized twice intramuscularly (i.m.) with 10^5^ PFU of extracellular *dl*5-29 virus or mock cell lysate in a 25 µl volume. Injections were carried out 21 days apart and with mice under isoflurane anesthesia.

### Behavioral tests

Breeding and infections were conducted in a separate space than the behavioral studies room. Mice were acclimated to the behavioral facility under a 12-hour light cycle (7a-7p) for at least 7 days prior to testing. Environmental conditions (test room lighting, temperature, and noise levels) were kept consistent and both male and female mice were included in the study. Singly-housed animals were excluded from behavioral assessment as isolation is known to alter behavioral task performance^89^.

### Novel object recognition

For the novel object recognition (NOR) test, animals were placed in a 30cm x 30cm open field containing two identical objects for 10 minutes^90^. Twenty-four hours later, mice were returned to the enclosure with one familiar object (from the previous day) and one novel object. Mouse movement in the enclosure was recorded (Canon VIXIA HF R800), and the amount of time mice spent with each object was quantified using open-source Matlab software that had been previously validated^91^.

Discrimination index (Time_novel_ – Time_familiar_)/Time_total_) was quantified for the 5-minute test trial to compare novelty preference while controlling for differences in individual mice’s locomotion. Unless otherwise indicated, the NOR test was conducted at 4-5 months post infection. All enclosures and objects were cleaned thoroughly with 70% ethanol between sessions to eliminate olfactory cues.

Inherent object preference was tested over a span of 5 minutes using naïve mice and object pairs were used only if no object preference was observed. Objects used in this study included: white plastic unicorn ducks, T-25 flasks filled with colorful aquarium pebbles, white plastic ducks, and pink plastic pig toys. Objects were balanced based on size and randomized for the familiar object training.

### Novel object location task

For the novel object location (NOL) task^45^, mice were placed in the same 30cm x 30cm enclosure for 10 minutes during the familiarization phase. However, instead of two different objects, mice were familiarized to two identical objects in two corners of the arena (example: horizontal in the top two corners of the arena). One hour later, mice were returned to the enclosure with the same identical objects, one moved to a novel location in the arena (example: diagonal with one object remaining in the top left and one object moved to the bottom right). Mouse interaction time was recorded for 5-minutes (Canon VIXIA HF R800) and the first 20 seconds of total object interaction time was used to compute the discrimination index for each mouse (see above). All analysis was done using Noldus Ethovision software^92^. All enclosures and objects were cleaned thoroughly with 70% ethanol between sessions to eliminate olfactory cues. Object locations were balanced between familiarization and testing phases. Objects used in this study: saltshakers filled with DRIERITE desiccant.

### Modified Barnes maze

The modified Barnes maze was performed on 5-month-old adult mice as previously described^53^. As illustrated in Figure 3, the maze consisted of a white 88.9 cm-diameter circle with 12 equally spaced holes around the circumference of a 15.24 cm-tall circular wall. Spatial cues consisting of large red letters (“W”, “S”, “T”, “O”) we positioned at 90-degree increments around the inside of maze. All holes aside from the escape hole were plugged. The arena was illuminated by fluorescent lamps to create an anxious environment. Mice were placed in the center of the platform facing the W and given 4 minutes to find the escape hole. If the mouse entered the escape hole before 4 minutes, the experiment ended. Mice that did not enter the escape hole were led to it by the experimenter and allowed to exit the escape hole. Animals exited the escape hole into their home cage to reinforce maze escape. Mice received a total of 3 trials per day with an inter-trial interval (ITI) of 1 hour. Training consisted of 4 consecutive days where mice learned to find and enter the training escape hole. Completion of training was followed by a 2-day rest period. The training escape hole location was then reinforced for 2 days of three trials each. Following 6 days of training, the escape hole was switched to the location directly opposite its training position and mice were run for two days of three trials each. Mouse movement in the training and reversal trials was recorded (Canon VIXIA HF R800) and analyzed using Noldus Ethovision^92^. Escape latency and distance traveled were quantified in a blinded fashion using Noldus Ethovision software, while total errors and errors per quadrant (nose-pokes into non-escape holes) were quantified manually by two blinded experimenters.

### Different Paired Associate Learning (dPAL)

dPAL was conducted using the Second Generation Bussey-Saksida Touch Screen Chambers for Mice as previously described^56,58,93^ (Lafeyette Instrument, model 80614A). Briefly, the testing was conducted in a trapezoidal operant chamber with a metal floor, a reward delivery machine, a touch screen, two infrared (IR) beams to detect mouse movement, and black Perspex sidewalls (see Fig. 4A). The chamber was housed inside a sound- and light-attenuating box to eliminate distraction from the outside environment. The chamber contained a house light, a tone generator, a ventilating fan, and an IR camera to monitor mouse performance throughout dPAL trials. The environment and data collection was controlled by ABET software provided by Campden Instruments. For the current experiment, a three-window (each window 7 cm x 7 cm) mask was placed over the touch screen to limit interaction zones. Liquid reward (evaporated milk, Nestle Carnation) was provided to motivate and reinforce task performance.

To further motivate mice to respond to a food reward, mice were food restricted and maintained at 85- 90% initial free-feeding body weight. Weights were taken daily for one week prior to food restriction and every day of the experiment to ensure mice maintained appropriate body weight. The mice were assigned a consistent touchscreen chamber throughout training and testing. Mice were initially habituated to the chambers with a 20-minute operant chamber session. The next day mice were habituated to an operant chamber for 40 minutes. For all following training sessions, mice were rewarded with evaporated milk (Nestle Carnation). Mice were trained to interact with the touchscreen through four serial training stages: “Initial Touch”, “Must Touch”, “Must Initiate”, and “Punish Incorrect”. During “Initial Touch” mice were presented with a white square stimulus in a pseudo-random window and trained to touch anywhere on the chamber touchscreen to receive a reward (criterion: completion of 30 trials within 60 minutes). If, during “Initial Touch”, mice touched the monitor where the white stimulus is presented, the mouse was rewarded with an excess of 3x the reward. During “Must Touch”, Animals were trained to nose-poke only the window where the white square stimulus was presented (criterion: completion of 30 trials within 60 minutes). In “Must Initiate,” mice were trained to break the IR beam directly above the reward magazine to initiate a trial and touch the presented stimulus to complete 30 trials in under 60 minutes. Finally, in “Punish Incorrect,” the final stage of training, if a window that did not contain the stimuli was nose-poked, animals were presented with a 5 second ITI signaled by house-light illumination and an incorrect tone. Following the 5 second ITI, mice underwent correction trials with the stimulus presented in the same window until a correct response was recorded. Collection of the reward in the magazine was followed by a 15 second ITI. Mice were required to achieve at least 80% correct in 30 trials within 60 minutes for two consecutive days before graduating to dPAL testing.

The following dPAL protocol for mice was based on the work of John Talpos et al.^58^, using the “Flower- Plane-Spider” stimulus combination shown in Figure 4. In dPAL, a stimulus, such as Flower, appeared in its correct touchscreen location (S+). A second stimuli, such as Plane, appeared in the incorrect location (S-). A dPAL trial began when the mouse nose-poked the illuminated reward chamber. This deactivated the reward light and triggered the display of the S+ and S- stimuli on the monitor.

Response to S+ triggered a reward tone, illuminated the reward chamber, delivered milk reward, and the removed visual stimuli from the screen. Collection of reward caused the reward chamber light to deactivate and initiated a 10s ITI. After the ITI, mice were allowed to initiate the next trial by nose-poking the illuminated reward chamber. Response to S- triggered a 10s time-out when stimuli was removed from the screen, illuminated the house-light for 5s, and cued an incorrect tone to play. After the ITI, the reward tray light was illuminated and mice nose-poked to initiate a correction trial.

Correction trials were not counted towards the total trials completed or the percent correct performance. During correction trials, mice were presented with S+ and S- in the same windows as during the incorrect trial. Mice repeated correction trials until the they picked S+ and proceeded to the next dPAL trial. The session finished after either 36 trials were completed, or 60 minutes had passed. Mice completed one testing session per day. Data was binned into 3-session blocks representing 108 trials each, completed across 3 sessions.

### IgG Purification and Delivery

IgG purification was performed on AKTA FPLC system via Protein G Agarose (Thermo Scientific Pierce) affinity chromatography as previously described^38^. Briefly, 5 ml murine serum was equilibrated (1:2) with 100 mM sodium acetate, pH 5, before the sample was applied to the column with a flow rate of 0.250 ml/min. The protein elution was achieved in a 15 ml volume with 100 mM glycine, pH 2.7, and quickly neutralized with 100 mM Tris-HCl, 0.5 M NaCl, pH 8.5. Elution fractions were then concentrated, buffer exchanged to PBS and filter sterilized. Purity was assessed via protein gel electrophoresis and concentration was determined via Nanodrop spectrophotometer. Dams were treated intraperitoneally (i.p.) with 1 mg of naïve or HSV immune IgG 3-5 days prior to parturition.

### Immunofluorescence

For antibody staining, brains were collected from PBS perfused mice, fixed with 4% paraformaldehyde, and cryopreserved in a sucrose gradient of 15%-30% at 4⁰C until they equilibrated and sank. Brains were then embedded in Tissue-Tek OCT compound (Sakura) and stored at -80⁰C prior to cutting.

Slices were prepared using Leica CM1860 UV cryostat at -15-20⁰C. Tissue was mounted onto Colorfrost Plus microscope slides (Fisher Scientific) and incubated with 0.01% BSA and 0.25% Triton X-100 for 20 minutes, then 5% normal goat serum and 0.25% Triton X-100 for 30 minutes for permeabilization and blocking. Primary and secondary antibodies were suspended in 1% NGS and 0.1% Triton X-100. Antibodies used in this study include rabbit anti-HSV gC (DAKO), Alexa Fluor 647 Dk anti-rabbit (Jackson ImmunoResearch), and 4’,6-diamidino-2-phenylindole (DAPI; Thermo Fischer Scientific; 1:5,000).

Microscopy images were acquired on a Zeiss Axio Observer.Z1 microscope and images shown are representative of at least three experiments. All image analysis was conducted using ImageJ software.

### Viral DNA extraction and HSV-1 genome copy number quantification

To quantify HSV-1 viral DNA (vDNA) in the brain of infected mice, we assayed the midbrain (MB) region of pups infected with 100, 500, and 1000 PFU at 5 dpi. MBs from mock-infected pups were used as a negative control. Genomic DNA from each individual MB was isolated via proteinase K, phenol-chloroform extraction, followed by ethanol precipitation. Following successful isolation of whole genome DNA, the copy number of HSV-1 per MB was determined using primers for VP16 (UL48; forward, 5’- AACCACATTCGCGAGCACCTTAAC-3’; Reverse, 5’-CAACTTCGCCCGAATCAACACCAT-3’. qPCR reactions were performed using the following components to a final reaction volume of 20 μl: 1X Luna Universal qPCR Master Mix (New England Biolabs), forward and reverse primers at a final concentration of 0.25 μM, 100 ng of DNA, and 5% acetamide. The qPCR reaction settings were as follows: 1 cycle at 95⁰C for 60 s, 42 cycles at 95⁰C for 15 s and 60⁰C for 30 s, followed by a melt curve with a range of 60-95⁰C. To determine HSV copy number, a standard curve was generated by diluting purified HSV-1 st17 genome into background mouse whole genome DNA isolate in 10-fold dilutions ranging from 10^6^-10^1^ copies. These dilutions were used to determine the total genome copy number in the isolated MB samples. To normalize the quantification of VP16, qPCR of a single-copy mouse gene, Adipsin, was performed. The mouse adipsin primer set used was: 5’-AGTGTGCGGGGATGCAGT-3’ (Forward), and 5’- ACGCGAGAGCCCCAGGTA-3’. A standard curve for adipsin was also generated by making standards of uninfected mouse DNA isolate that range from 10^6^-10^1^ copies. Based on the qPCR amplification of both VP16 and adipsin in test samples, HSV-1 copy number was calculated as described previously^67^.

### Statistical analysis

Prism 10 (GraphPad) software was used for statistical tests. For survival studies, nHSV-infected mice were compared to mock-infected controls using the Log rank Mantel-Cox test to determine p-values. For copy number analysis, HSV dosage groups were compared to each other via a two-way ANOVA with Bonferoni’s test for multiple comparison. For NOR, NOL, and MBM performance, groups were compared using unpaired t-tests. For dPAL, groups and time points were compared to each other via two-way mixed ANOVA, with Tukey’s test for multiple comparisons to determine p-values.

## Acknowledgments

We thank all members of the Leib and Nautiyal Labs, Margaret Ackerman, Pamela Rosato, Alex Skorput, Kirk Maurer, David Knipe, Bryan Luikart, Steve Fiering, Charles Sentman, Hermes Yeh, the late David Bucci, and the Jones Media Center for materials and/or helpful discussion. Graphical figures were created with BioRender.com.

## Author contributions

D.A.L., C.D.P., and A.D. designed the research with advisement from K.M.N. C.D.P., A.D., S.A.T., C.G., E.V., J.M., and R.A.V. performed the experiments. A.D., C.P., and D.A.L. wrote the manuscript with the assistance of other authors.

